# Automated Quality Control of Time-Course Imaging from 3D in vitro cultures

**DOI:** 10.1101/2025.03.31.646437

**Authors:** Eric Cramer, Tamara Lopez-Vidal, Jeanette Johnson, Vania Wang, Daniel Bergman, Ashani Weeraratna, Richard Burkhart, Elana J. Fertig, Jacquelyn W. Zimmerman, Laura M. Heiser, Young Hwan Chang

## Abstract

Longitudinal imaging of 3D cell cultures like tumor organoids and spheroids offers crucial insights into cancer progression and treatment. However, spatial displacement during time-course imaging, caused by matrix detachment or experimental artifacts, can confound analyses. Existing computational methods struggle to address this issue. We present a new algorithm to evaluate data integrity and rectify mislabeling in longitudinal imaging of 3D cell culture. Our algorithm integrates permutation-based optimization with Procrustes analysis. By using X and Y coordinates of images, it accurately reorders, matches, and aligns object positions across time points, correcting for rotation, translation, and small movements. Validation with simulated data confirmed its accuracy and robustness. Applied to longitudinal imaging of tumor spheroids, our algorithm revealed frequent displacement amongst the spheroids between time points and corrected many mislabeled images. This computationally efficient and adaptable method needs no experimental adjustments and presents a readily accessible solution for data quality control.

**Motivation:** Three-dimensional (3D) in vitro models, such as tumor organoids and spheroids embedded in an extracellular matrix, are increasingly vital for studying normal and disease biology, including drug responses.^1–3^ A key advantage of these models is that imaging platforms can perform continuous longitudinal imaging to track phenotypic changes. However, common issues in 3D techniques, such as matrix shifts during experimental setup or image capture, can introduce technical artifacts that affect downstream analyses. Currently, no automated analytical approaches exist for assessing or correcting technical artifacts. Here, we introduce a robust, automated algorithm for assessing the quality of time-course image data and, in some cases, correcting object mislabeling to enable accurate tracking of individual spheroids over time. This approach relies only on image metadata, requiring no experimental modifications. It offers a readily implementable solution for improving data integrity and reproducibility and enhancing the reliability of longitudinal 3D cell culture studies.

## Introduction

Three-dimensional (3D) cell culture models like tumor spheroids are gaining traction in cancer research for their ability to recapitulate key aspects of in vivo tumor biology – capturing spatial architecture, cellular heterogeneity, cell-cell interactions, and drug resistance mechanisms better than traditional two-dimensional (2D) monolayer cultures.^1–5^ Longitudinal imaging with bright field, phase-contrast, and fluorescence microscopy enables better monitoring of tumor spheroid growth, morphology, specific cellular processes, and responses to experimental conditions.^1,6,7^ Proper implementation enables single-cell and region-of-interest (ROI) analyses, revealing dynamic changes and spatial reconfiguration of cell types in response to external stimuli.^8^ These approaches enable high-throughput drug screening, capturing factors like drug penetration, efficacy, toxicity, and microenvironmental factors such as cell-cell interactions and the development of hypoxic regions.^1,3,9,10^ This makes longitudinal monitoring of tumor spheroids a valuable tool for preclinical research and therapeutic evaluation.^1,11^

A key challenge in time-series imaging of tumor spheroids is spatial displacement caused by experimental artifacts and technical effects.^1,6^ Long-term cultures, repeated media changes, or mechanical disturbances can shift an embedded object’s position, particularly if a well plate insert or the collagen matrix detaches (Fig. 1A and 1B). If detected, a displaced matrix will require repeating experiments, wasting time and resources. If not noticed, such shifts result in mislabeling sequential images, making it difficult to track the dynamic behaviors of individual spheroids accurately over time (Fig. 1C and 1D).^10,11^ These displacements can confound downstream analyses, leading to misinterpretation of growth dynamics and treatment responses.^8^ Therefore, new computational techniques are needed to quantify when such errors in longitudinal imaging have occurred. When detected, this can indicate when experiments need to be repeated. Further computational techniques to correct these artifacts can enable use of these experiments despite technical artifacts, reducing the wasted time and resources from redoing experiments.

**Figure 1.**
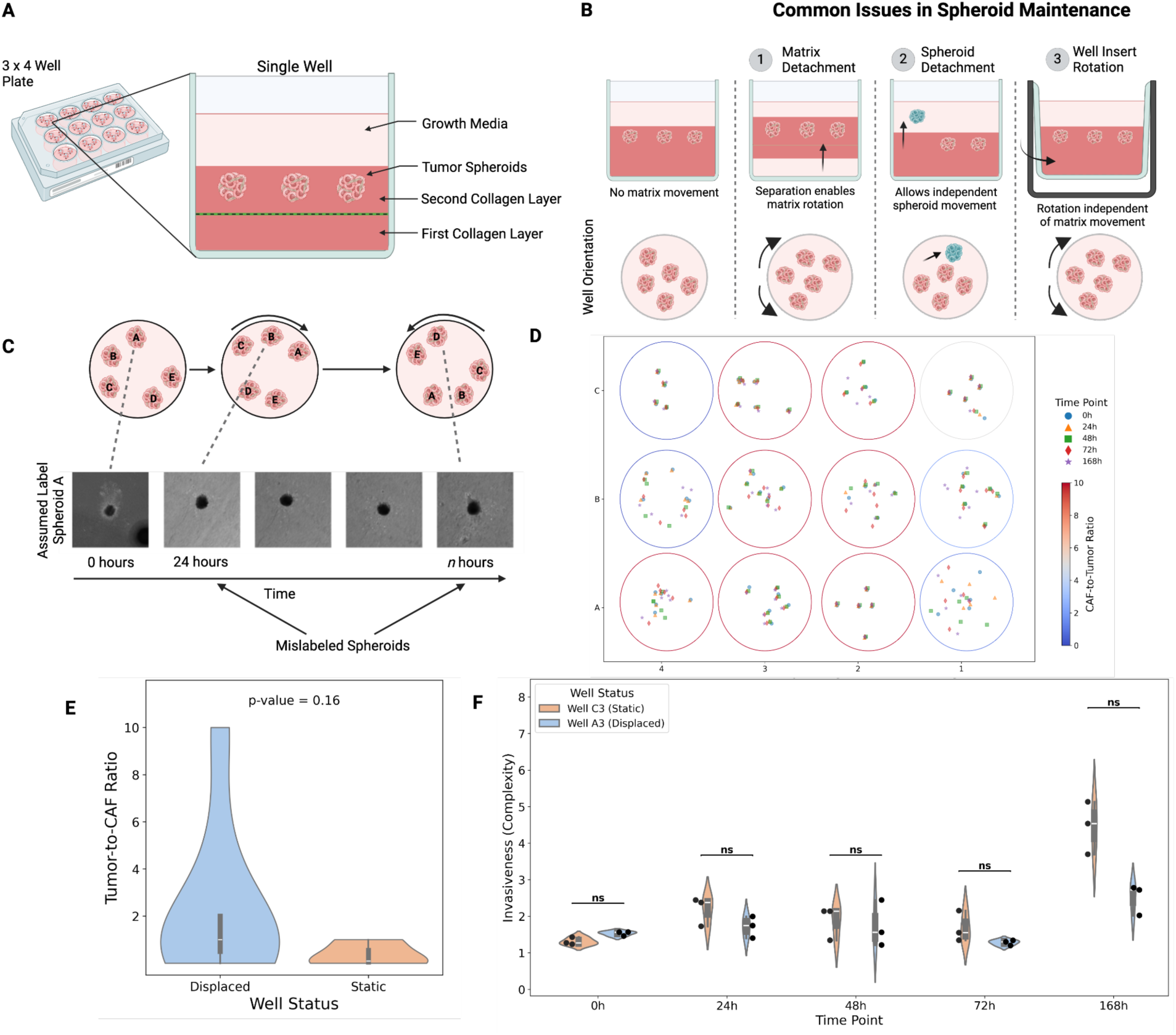
Experimental Setup and the Effect of Collagen Detachment on Spheroid Mislabeling. **A)** Schematic of a well plate and a single well, illustrating the layered structure of the spheroid culture system (first collagen layer, second collagen layer, tumor spheroids embedded within collagen, and growth media). **B)** Illustration of scenarios where recording spheroid measurements may become mislabeled over time. Change in spheroid positions due to rotating matrix or well plate inserts, or displacement of a single spheroid between time points can lead to recording incorrect measurements of each spheroid over time. **C)** Illustration of how well rotation leads to mislabeling measurements of a spheroid over time. The assumed labels based on initial positions do not match the actual positions of the spheroids. Brightfield images illustrate this mislabeling. **D)** Schematic of the biological dataset, showing a 3×4 well plate containing tumor spheroids with varying cell type ratios. Each circle represents a well, with spheroid locations at different time points indicated by color and shape. **E)** Violin plot showing the relationship between the ratio of Panc 10.05 tumor cells to hT231 fibroblast cells comprising the spheroids in each well of the well plate to the status of the well as exhibiting rotation or spheroid displacement. Comparison by the Kruskall-Wallis test did not show a significant difference in tumor-to-CAF ratio between displaced and static wells. **F)** Invasiveness (quantified by complexity) of spheroids over time, categorized by whether the well exhibited displacement of the spheroids or not (“well status”). Comparison by the Kruskall-Wallis test did not show significant differences in complexity at any time point.

Current computational approaches struggle to quantify drift and correctly label 3D cultures in longitudinal imaging due to the dynamic and evolving nature of spheroids over time. Matching images to the label associated with the nearest neighbor between time points is unreliable because object shape, texture, and other characteristics change, altering their appearance as they grow and invade the surrounding matrix.^7,12,13^ Low-dimensional feature representations, such as latent embedding from autoencoders, are also limited because they do not incorporate the global positional information needed to track objects across time points.^14^ Traditional image registration techniques, which rely on fixed spatial landmarks, are also ineffective since embedded objects typically lack stable reference points, particularly when imaged as isolated objects rather than within a larger well plate context (Fig 1C).^15^ Furthermore, in the absence of a large scan image capturing the full well plate at each time point, object-based landmark registration becomes infeasible. As a result, current approaches fail to provide a robust solution for detecting and correcting matrix displacement, increasing the likelihood of data misinterpretation and experimental artifacts.

To overcome these challenges, we propose an automated algorithm that ensures accurate spheroid-matching by leveraging positional geometry across time points. Using a dataset of tumor spheroids embedded in a collagen matrix and imaged every 24 hours, we demonstrate the effectiveness of our computational algorithm in detecting and correcting matrix displacement. Our approach utilizes image metadata-derived coordinates to construct a polygon representing spheroid positions within the well. By applying an affine transformation, our algorithm corrects for rotational, translational, and small movements in spheroid positioning.

This approach enables precise tracking of individual spheroids over time, ensuring data integrity without requiring additional experimental modifications. By implementing this geometric alignment framework, we provide a robust quality control step for longitudinal 3D imaging, enhancing the reliability and interpretability of spheroid-based assays in cancer research.

## Results

### Spheroid displacement was not associated with biological conditions

To investigate spheroid behavior over time, we generated tumor spheroids with varying cancer associated fibroblast (CAF) to tumor cell ratios, embedded them in a collagen mixture, and replated them into pre-coated wells (Fig. 1A & Fig. 1D). We ensured that each well contained four to six replicates of each cell ratio (see Methods Spheroid Generation) within the collagen matrix. Each spheroid was then imaged at 24, 48, 72, and 168 hours to monitor their growth and invasion while assessing the integrity of the surrounding matrix. Common issues in long-term imaging experiments include matrix shifting and the rotation of well plate inserts, which can displace spheroids within a well relative to their position at the initial time point (Fig. 1B).^9^ Without global positional information for the entire well plate at each time point, these shifts can cause misattribution of spheroid identities as seen in Figure 1C, confounding the interpretation of a single spheroid’s dynamics and leading to erroneous conclusions. Following the image position acquisition methods, we obtained a table of XYZ coordinates for all spheroids at each time point. Analysis of the positional data revealed that in 9 out of 12 wells, at least one spheroid exhibited substantial displacement at some point during the experiment (Fig. 1D). This finding highlights the prevalence of positional shifts in long-term spheroid culture experiments, necessitating a correction. Thus, we categorized the status of wells that featured spheroids with displacement as “displaced” and those that did not feature spheroid displacement as “static.”

To determine whether cell type ratio influenced spheroid displacement, we assessed the relationship between tumor-to-CAF ratio and well status via the Kruskall-Wallis test (Fig. 1E). The results showed no significant association between spheroid displacement and the tumor-to-CAF ratio of the spheroid, indicating that the experimental variable did not drive spheroid displacement. Next, we investigated the potential impact of well status on spheroid invasiveness (the experimental outcome measure of interest), measured with complexity– a shape descriptor calculated from each spheroid’s segmentation mask. Higher complexity values indicate greater invasiveness, as spheroids develop irregular edges and spread into the matrix.^12^ Violin plots (Fig. 1F) depict the distribution of complexity values over time for spheroids composed of a 1:1 tumor-to-CAF ratio in two example wells: one where spheroid coordinates remained static and another where displacement occurred. No statistically significant differences in complexity were observed between the two groups at any time point, suggesting that matrix shifts did not alter the biological conditions of the experiment and impact the overall invasiveness of the spheroids.

Since spheroid complexity remained consistent across well status over time, we inferred that their morphological behavior—and thus invasiveness—was not affected by displacements. This stability allowed us to proceed with analyzing individual spheroid dynamics. To do so, we first needed to correct mislabeling of observations across time points by developing an algorithm to accurately pair each image with its corresponding spheroid.

### A two-step algorithm for matching spheroid images from longitudinal imaging

To correct errors in image assignments caused by spheroid displacement, we developed a custom algorithm that automatically reassigns mislabeled images to the correct spheroid at each time point (see Methods section on Image Matching; Fig. 2A). The X and Y coordinates of each spheroid’s image are first extracted from that image’s metadata (see Methods section on Image Position Acquisition). The first step of the algorithm establishes a reference configuration of the spheroids at the initial time point (t=0). A permutation-based optimization strategy then considers all possible orderings of spheroids and selects the permutation that minimizes the L2 norm between the pairwise Euclidean distance matrix of the current configuration and the reference matrix at t=0.

**Figure 2.**
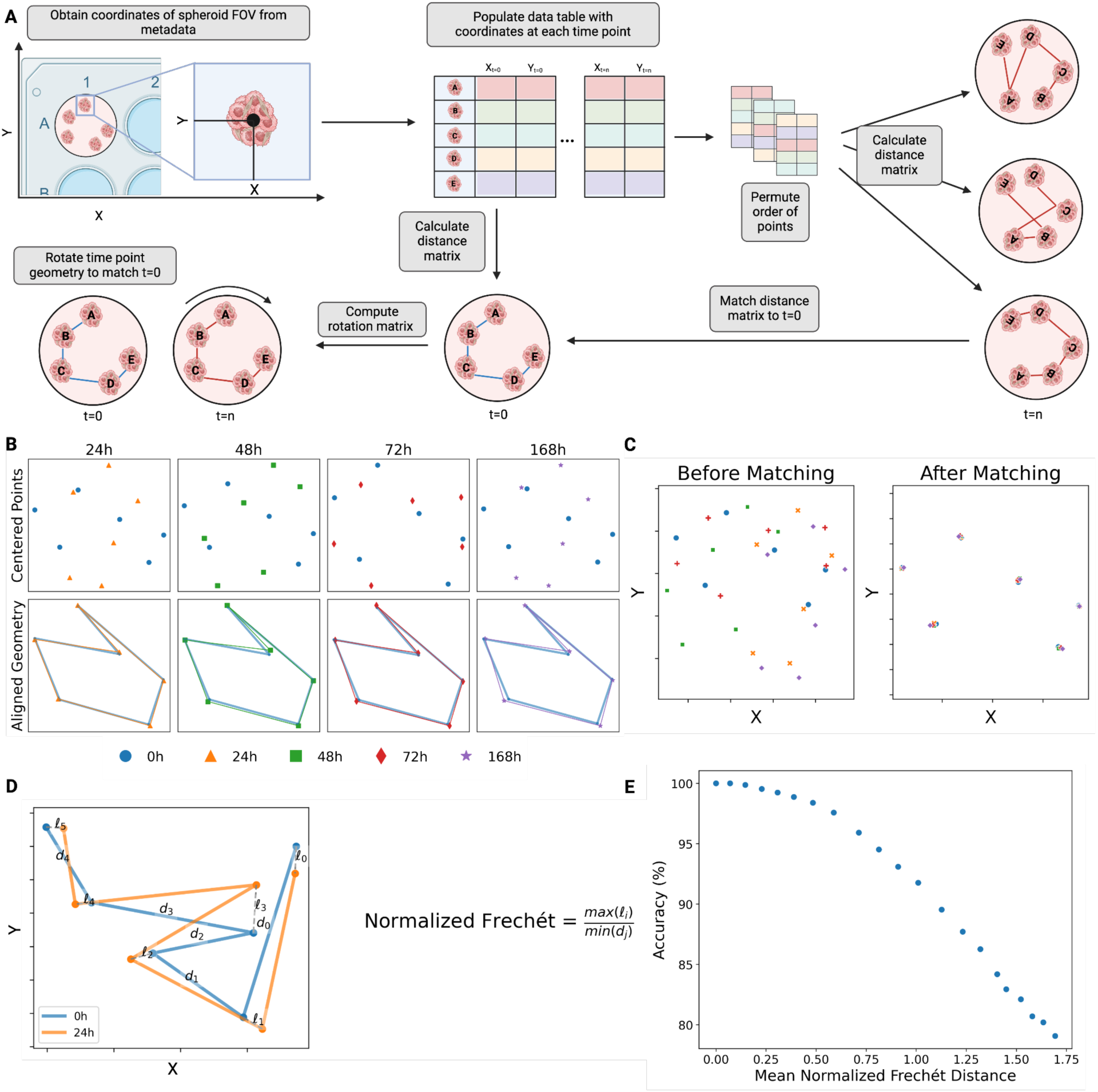
Spheroid Alignment Algorithm and Evaluation of Performance. **A)** Schematic of the spheroid alignment algorithm. Spheroid coordinates are extracted from image metadata. Pairwise distances between spheroids at t = 0 are calculated and serve as a reference. For subsequent time points, a permutation-based optimization strategy identifies the spheroid ordering that minimizes the difference between pairwise distance matrices relative to the reference. Procrustes analysis is then used to determine the optimal rotation and translation to align the reordered spheroids to the t = 0 positions. **B)** Simulated spheroid locations (n=5 spheroids) at different time points (t = 0, 24, 48, 72, and 168h) before and after alignment. Lines connect spheroids to visualize the geometric transformations. Shared legend with Figure 1C. **C)** Alignment results for the simulated data. Left: Spheroid positions before alignment. Right: Spheroid positions after alignment to the t = 0 reference. **D)** Illustration of the normalized Fréchet distance calculation. The Fréchet distance quantifies the similarity between two curves (here, the paths connecting spheroids in a specific order). Normalization by the minimum nearest neighbor distance adjusts for variability in scale between different samples. **E)** The relationship between the normalized Frechét distance and the matching algorithm’s accuracy calculated from a data set of 105,000 simulated wells featuring increasing amounts of single-spheroid positional perturbations. A normalized Frechét distance of 1.0 corresponds to approximately 90% accuracy.

The second step of the algorithm is an alignment process that applies Procrustes analysis to the optimal permutation of the spheroids identified during the first step. Procrustes analysis is a statistical method to compare shapes by minimizing the sum of squared distances between corresponding points and transforming them into a state of maximal superimposition. This technique finds the optimal rotation, translation, and scaling to align two or more sets of data points. In our implementation, the coordinates of the reference configuration and the optimal permuted configuration are first centered by subtracting each configuration’s centroid. The singular value decomposition (SVD) of the centered coordinates is then used to calculate a rotation matrix and translation vector. The rotation matrix and translation vector are then applied to the centered coordinates of the optimal permuted configuration to align the permuted configuration to the reference. The optimal permutation of the points is then used to re-order the raw coordinates to correspond to their aligned counterparts and assign correct labels to each image.

### Development of an automated quality-control and accuracy metric using simulation studies

To evaluate the algorithm’s performance, we first tested it on synthetic data, simulating the displacement of six spheroids under randomized spatial arrangements. Each simulation introduced random rotations, translations, and positional jittering, mimicking experimental disturbances. Figure 2B depicts the geometry matching process for an example simulation, with the initial unregistered spheroid positions at various time points and the resulting alignment of the algorithm, which rotates, translates, and stretches the connections between the simulated points to match their geometries to their configuration at the initial time point. As shown in Figure 2C, the algorithm successfully matched spheroids at each time point, restoring their spatial relationships to the initial baseline configuration.

To quantitatively assess the performance of the algorithm, we adapted the Fréchet distance, which measures the smallest maximum distance necessary to connect two paths of points in space while accounting for the ordering of the points.^16^ To allow comparisons across different spatial scales, we normalized the Frechét distance by dividing it by the minimum initial distance between any two spheroids (Fig. 2D). We chose to normalize by the smallest distance because it represents a conservative estimate for the threshold at which a spheroid’s nearest neighbors would change in the squared Euclidean distance matrix, and subsequently compromise the permutation strategy in the first step of our algorithm.

To further evaluate the algorithm’s robustness, we simulated displacement of a single spheroid at varying distances from its original position, ranging from 0% to 200% of the minimum nearest-neighbor distance, increasing in 10% increments. Each perturbation level was tested across 100 independent simulations, and the algorithm was applied to 50 randomly generated initial spheroid configurations. The algorithm’s accuracy was determined based on the percentage of correctly matched spheroids for each simulation. For each perturbation level, the average accuracy was plotted against the average normalized Frechét distance to establish a relationship between our performance metric and algorithm accuracy (Fig. 2E). The results showed that a normalized Frechét distance of 1.0 corresponded to approximately 90% accuracy, establishing a threshold for extreme perturbations. Notably, this metric can also be used for automated decision-making by the algorithm, eliminating the need for manual visual evaluation of the geometry graph (Fig. 2B). The algorithm effectively corrected for rotation, translational, and random shifts within the range of typical experimental shifts.

### Matching algorithm identifies and corrects displacement errors in experimental data

After validating the robustness of our algorithm on simulated data, we applied our algorithm to each well in the plate from our data set of tumor spheroids with varying tumor-to-CAF ratios, imaged over 168 hours (Fig. 1D, Supplemental Fig. 1 & Supplemental Fig. 2). Representative examples in Figure 3A of wells from the well plate illustrate the algorithm’s performance across a range of scenarios; examples include successful matching in wells with varying displacement level (Well A1), partial success, where certain time points were accurately matched while others were poorly matched (Well B1), a case where matching failed due to extreme displacement or confounding factors (Well B4), and a case where a single spheroid’s coordinates shifted over time (Well C1). These examples demonstrate the variability of scenarios in which the algorithm was able to successfully pair spheroids with their correct associated images. An example of manual visual inspection for a displaced well is shown in Supplementary Figure 3.

**Figure 3.**
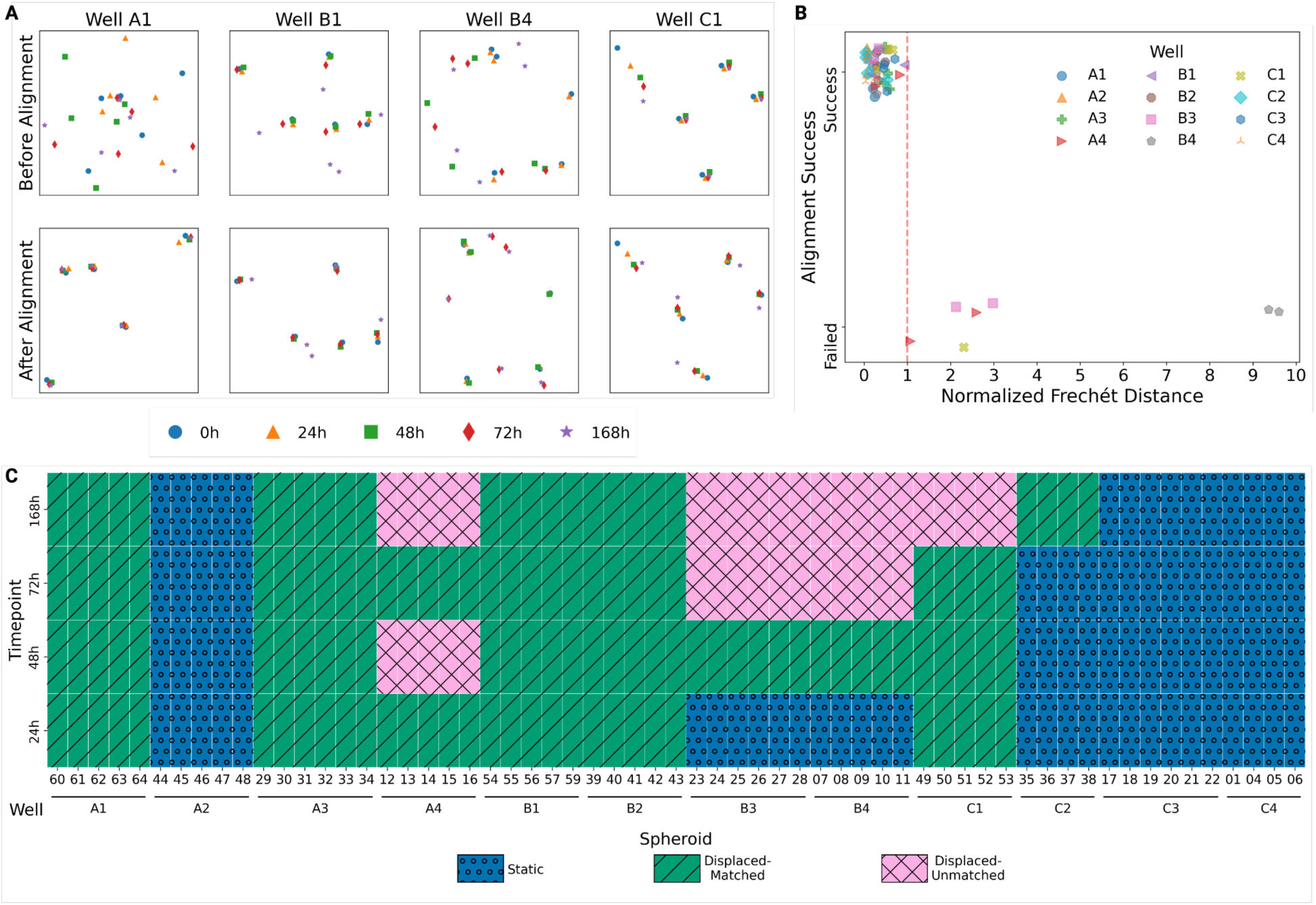
Performance of the Matching Algorithm on Biological Spheroid Data. **A)** Example results for selected wells, showcasing scenarios of successful matching, partial matching, matching failure, and the impact of when a single spheroid’s position changes over time. Each panel displays spheroid positions before and after the matching algorithm. **B)** Scatter plot showing the relationship between the normalized Frechét distance measured after matching at each time point for each well and the success or failure of that match. Dashed line at Normalized Frechét Distance of 1.0 indicates cutoff for algorithm alignment success determined during simulation studies. **C)** Heatmap summarizing the status of each spheroid: static (no displacement and no correction needed), recovered (successfully matched), and unrecovered (failure to match). Algorithm success relative to time point is recorded on the Y-axis.

To quantify the success of matching, we calculated the normalized Frechét distance for each well at each time point and compared it between successfully and unsuccessfully matched wells (Fig. 3B). Wells with normalized Frechét distances below 1.0 consistently exhibited successful matching. This supports our simulation findings that extreme positional shifts reduce algorithm accuracy (Fig. 2E). Furthermore, this finding established that our normalized Frechét distance can serve as a quantitative quality control measure of spheroid displacement over the course of experiments. Additionally, we observed that smaller nearest-neighbor distances at the initial time point (t=0) can cause increased normalized Frechét distances, suggesting that spheroids should not be embedded too proximally to each other to facilitate accurate matching (Fig 2B, Well B4).

To establish a comprehensive assessment of the algorithm’s ability to correctly label spheroids and recover the dynamics of individual spheroids across the experimental dataset, we calculated the matching success for each spheroid at every time point. We found that nine of the twelve wells demonstrated spheroid coordinate displacement during at least one time point, leading to uncertainty in 66% of the total measurements collected during the experiment. Our algorithm successfully recovered or validated 77% of potentially mislabeled spheroid measurements (Fig. 3C), demonstrating its ability to deconvolute confounded data and improve experimental reliability.

## Discussion

This study addresses a critical challenge in longitudinal imaging of tumor spheroids — mislabeling images of spheroids due to rotation and translation of matrices or well-plate inserts over time. Alternative experimental modification could be considered to ensure that images of tumor spheroids will be correctly labeled. For instance, incorporating well plate inserts with grids etched beneath the matrix or embedding fiducial markers could provide fixed reference points for tracking.^17^ Alternatively, acquiring large scan images of the entire well plate at each time point, in addition to individual fields of view, would allow for post-hoc reconstruction of spheroid trajectories even in the presence of matrix displacement. However, these alternatives introduce additional complexity or cost, such as new equipment or data storage requirements, to the experimental workflow.

Our matching algorithm provides a robust, flexible, and computationally efficient solution that ensures accurate tracking of tumor spheroids across time points. In addition, our development of the normalized Frechét distance provides a quantitative metric for automated quality control of tumor spheroid experiments as well as evaluating algorithm success. By calculating the normalized Frechét distance after alignment, researchers can determine the degree of change experienced by their spheroids over the course of an experiment as well as evaluate algorithm success.

Another key strength of this approach is its adaptability to existing experimental workflows. As it relies solely on field-of-view coordinates extracted from imaging metadata, our algorithm is broadly applicable to different types of assays that use longitudinal imaging of 3D cultures such as organoids embedded in extracellular matrix. The implications of this matching algorithm therefore extend beyond the specific context of spheroid tracking presented in this study. By relying solely on image coordinates, the algorithm may be applied to data generated by a wide range of experimental protocols and imaging platforms and be used prospectively to ensure data integrity in new studies or retrospectively to analyze existing datasets.

### Limitations of the study

However, the study does have limitations. The first limitation is that the biological data set with which we validate the algorithm is small. This may limit inference of potential biological findings in this study, but it does not detract from the demonstration of the algorithm itself. The second limitation is intrinsic to our algorithm as its performance depends on the initial spatial arrangement of objects. If objects are positioned in a perfectly symmetrical polygonal configuration (e.g., a square or regular hexagon), rotational symmetry can lead to ambiguities in alignment, making it difficult to correctly permute objects across time points. Additionally, matrix degradation or fragmentation over time can introduce inconsistencies, further challenging the ability to maintain object correspondence. Importantly, however, in cases of extreme geometric misalignment during the second step of the algorithm or matrix degradation, algorithm failure itself may serve as a useful quality control step, indicating potential structural instability within the matrix and highlighting concerns about the experimental setup.

Despite these limitations, our proposed algorithm and quality control metric present an automated and unbiased technique to ensure that the observed phenotypes of tumor spheroids embedded in collagen matrices are not related to an experimental artifact. By ensuring data integrity, the algorithm can prevent erroneous conclusions drawn from mislabeled data. This is critical for studies looking at the dynamic behaviors or spatial arrangements of single cells within regions of interest of a matrix-embedded object.^8,12,18^ Furthermore, provided measurements of biological interest remain consistent across displaced and static conditions, our approach offers the opportunity to recover previously unusable data affected by matrix displacement. This would maximize the value of existing experiments. The core principles of the algorithm—permutation optimization combined with Procrustes alignment—can be readily adapted to other applications involving spatial misalignment, such as organoid invasion analysis or tracking tissue architecture landmarks across adjacent sections.^19,20^

## Conclusions

This study introduces a novel matching algorithm to correct object mislabeling caused by matrix displacement or detachment in long-term culture experiments. Spatial misalignment presents a significant challenge for accurately tracking embedded objects and quantifying dynamic processes such as a tumor spheroid invasion. Our algorithm provides a robust and flexible solution by combining a permutation-based optimization strategy with Procrustes analysis, enabling precise matching of object positions across multiple time points. This method offers a valuable tool for ensuring data integrity and improving the accuracy of downstream analyses in assays using 3D tissue culture embedded in a matrix. Importantly, it achieves this without requiring modifications to experimental protocols, making it an efficient and scalable solution for researchers. The adaptability and computational efficiency of this algorithm make it suitable for a wide range of applications beyond spheroid tracking, wherever correcting spatial misalignment is crucial for accurate data interpretation.

## Methods

### Cell Line Culturing

The human CAF cell line hT231 was generated by digesting a resected pancreatic tissue sample following standard organoid line establishment protocols.^21^ The undigested fibrous tissue was then placed into cell culture plates with DMEM supplemented with 10% fetal bovine serum (FBS, Thermo Fisher Scientific) and penicillin-streptomycin (Gibco, 5140122). The 2D cells that proliferated were then immortalized by viral transduction with the SV40 LT-pLVX-SV40 LT-IRES-tdTomato plasmid, as previously described.^22^ The hT231 cell line and a pancreatic tumor cell line Panc10.05 and were cultured in RPMI 1640 Medium with L-Glutamine (RPMI, Thermo Fisher Scientific, 11-875-085) with 10% FBS, 0.1% Amphotericin B (Sigma, A2942), and 1% penicillin-streptomycin.^23^ Both cell types were Mycoplasma tested using Invivogen MycoStrips (rep-mys-100).

### Staining

To differentiate cells, CytoPainter Cell Proliferation Staining Reagents (Abcam) were used to stain each cell type. Panc10.05 was labeled for green fluorescence (ab176735), and HT231 was labeled for red fluorescence (ab176736), following the manufacturer’s instructions.

### Spheroid Generation

A 96-well plate (Celltreat, 229576) was treated with 50 µl of 1.5% Invitrogen UltraPure Agarose (#16500100) and allowed to solidify for 1 hour at room temperature. Panc 10.05 cells were mixed with hT231 cells at 1:0, 1:1, 2:1, 5:1, 10:1, 1:2, 1:5 and 1:10, tumor cell-to-CAF ratios. For each ratio, 4-6 technical replicates of 10,000 cells/well were seeded in a final volume of 200 µL/well of culturing media and onto the 1.5% agar-treated plate. Plates were maintained at 37 °C 5% CO2 for 4 days until spheroid formation was microscopically observed using the Nikon ECLIPSE Ti2 inverted microscope.

The spheroids were resuspended in 300 µl of Collagen Mix, which consists of 37% Rat Tail Collagen 1, 37% PBS (Corning, 21-040-CV), 10% FBS, 11.3% EMEM Media (Quality Biological, 112-018-101), 0.8% L-glutamine, and 5% NaHCO3 (Quality Biological, 118-085-721).^7^ Each spheroid ratio was then replated into a well of a 24-well SensoPlate™ (Greiner Bio-One, 662892), which had been pre-coated with 300 µl of the same Collagen Mix. Each well contained six spheroid replicates with identical ratios of Panc 10.05 and hT231 cells distributed within the collagen matrix.

### Image Position Acquisition

To visualize spheroid invasion, imaging was carried out with a Nikon ECLIPSE Ti2 inverted microscope using 10x and 20x objectives. Spheroids were imaged at 0, 24, 48, 72, and 168 hours after embedding. For each spheroid, five Z-stack images were taken across a 60 µm range using both brightfield and fluorescent imaging, with fluorescence captured using 647 nm and 488 nm lasers. The cell culture plates were placed on a live cell stage and maintained at 37°C with 5% CO₂ during imaging. At the initial time point (t=0), a large scan image of the entire well plate was captured, and spheroid positions were manually selected and saved as an XY Multipoint file. For each subsequent time point, the multipoint file from the initial time point was loaded, each spheroid was centered, and the Z-plane was refocused while keeping the PFS (Perfect Focus System) active. This ensured that all spheroid locations were imaged sequentially, reducing the need for manual adjustments. The collected spheroid images were then saved in a Nikon multipoint file (.nd2) for that time point. For each time point’s ND2 file, the X and Y coordinate tables were extracted from the metadata and exported. Meanwhile, the multipoint file was split into single-frame files and stored in a separate directory for downstream analysis.

### Image Matching

A custom algorithm corrected for potential displacement of tumor spheroids within wells of the well plate by matching the coordinates of images at later time points to the coordinates of the image taken at the first time point. Using well boundary coordinates and the X and Y locations of individual spheroid images from the metadata of the multipoint files, spheroids were assigned to their respective wells at each time point (t = 0, 24, 48, 72, and 168 hours). Spheroid locations in each well at t = 0 served as the reference configuration. For each subsequent time point, spheroid coordinates were centered by subtracting the mean X and Y positions. Pairwise squared Euclidean distances between spheroids were calculated for both the reference and current time point configurations. The optimal ordering of spheroids at the current time point was identified by a permutation-based optimization strategy as follows. This strategy iteratively permuted the order of spheroids, calculated the pairwise squared Euclidean distance matrix for each permutation, and selected the permutation that minimized the induced L2 norm of the difference between the current and reference distance matrices (Fig. 2A). The induced L2 norm was chosen for this application because it would optimize for the smallest of pairwise distances between each set of coordinates without assumptions about the order of the points.^24^

Procrustes analysis, based on singular value decomposition (SVD), was then applied to determine the optimal rotation and translation aligning the reordered current time point configuration with the reference configuration. Procrustes analysis is a statistical method to compare shapes by minimizing the sum of squared distances between corresponding points and transforming them into a state of maximal superimposition.^25^ This technique finds the optimal rotation, translation, and scaling to align two or more sets of data points. The detailed steps of the algorithm, including the Procrustes analysis and permutation optimization, are outlined as pseudocode in Algorithm 1. Specifically, let *A* be an n x 2 matrix representing the centered reference configuration and *B* be an n x 2 matrix representing the centered, reordered current time point configuration. The algorithm proceeds by calculating the SVD of *B^T^A* and uses the resulting matrices to compute the optimal rotation and translation. This procedure robustly aligned spheroid locations across time, correcting for potential displacements and rotations of the spheroids within the well.

**Algorithm 1.**
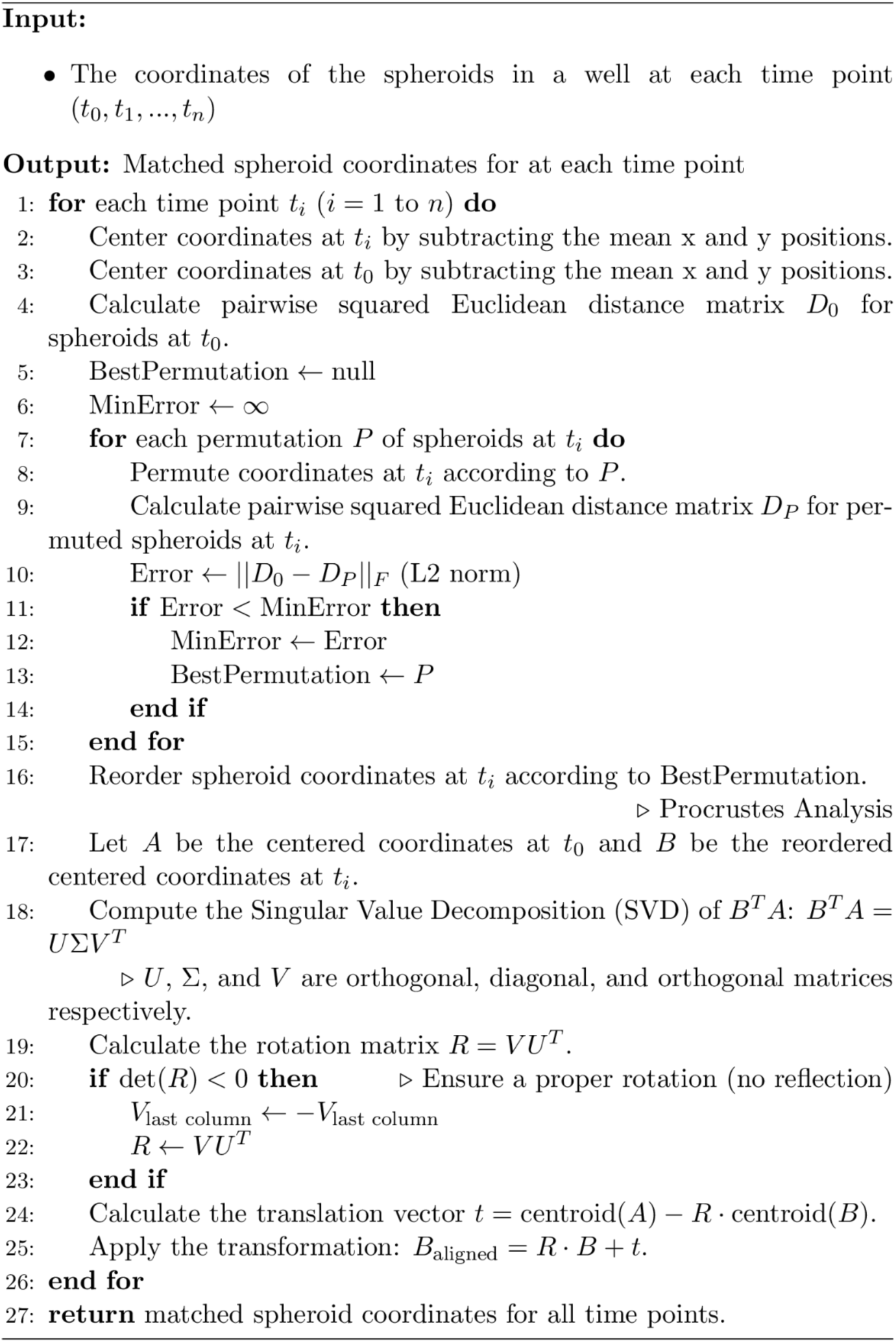
Spheroid Matching Algorithm.

### Quantifying Invasiveness

Each image for each spheroid was analyzed using a custom analysis pipeline implemented in Python version 3.10. The microSAM and napari packages were used to generate initial segmentation masks for each spheroid, and masks were post-processed to fill gaps and remove noise using the scikit-image and scipy packages.^26–28^ The complexity of the contour in each mask was analyzed to quantify the invasiveness of each spheroid at each point in time (t = 0, 24, 48, 72, and 168 hours).

### Statistical Analysis

After matching and image label correction, the trajectories of each spheroid’s invasiveness between the rotated and adhered conditions were compared via a non-parametric Kruskal-Wallis test using Python 3.10 and the scipy package.

## Resource Availability

### Lead Contact

- Requests for experimental protocols and reagents can be directed to Dr. Jacquelyn W. Zimmerman. Questions regarding the algorithm can be directed to Dr. Young Hwan Chang.

### Materials Availability

- This study did not generate new unique reagents.

### Data and Code Availability

- All software used in the study is detailed in the Methods section and supplementary material. All scripts are available via GitHub: https://github.com/emcramer/align_spheroid/releases/tag/v0.0.1-alpha
- All data generated in this study are available within the article and will be made available on Zenodo.
- Any additional information needed to reanalyze the data reported in this paper is available from the lead authors by request.

## Acknowledgment

We thank David Tuveson, Dennis Plenker, and Hardik Patel for kindly sharing the hT231 cell line, Dr. Elizabeth Jaffee for her mentorship, and feedback from Genevieve Stein-O’Brien and Paul Macklin. This work was supported by the Jayne Koskinas Ted Giovanis Foundation for Health and Policy grant awarded to L.H.; E.J.F was supported by NIH/NCI U24CA284156; A.W and V.W were supported by the T32CA153952; E.C. was supported by the NIH/NCI 1T32CA254888-01, and by the NIH/NIGMS 5T32GM141938-04. J.W.Z and T.L.V. report funding for the spheroid and CAF experiments from the Charles and Margaret Levin Family Foundation, and the Dana & Albert R. Broccoli Charitable Foundation. The research reported in this publication used computational infrastructure supported by the Office of Research Infrastructure Programs, Office of the Director, of the National Institutes of Health under Award Number S10OD034224. The content is solely the responsibility of the authors and does not necessarily represent the official views of the National Institutes of Health.

## Conflicts of Interest

EJF was on the scientific advisory board of Resistance Bio/Viosera Therapeutics, was a paid consultant for Mestag Therapeutics and Merck, and received grants from Roche/Genentech, Abbvie Inc, National Foundation for Cancer Research, and Break Through Cancer outside the scope of this work. J.W.Z. reports grant funding support (to Johns Hopkins) and travel from Roche/Genentech and funding from Break Through Cancer outside the submitted work, and honoraria from Sermo and ZoomRx.

## Supplemental Information

**Supplementary Figure 1.**
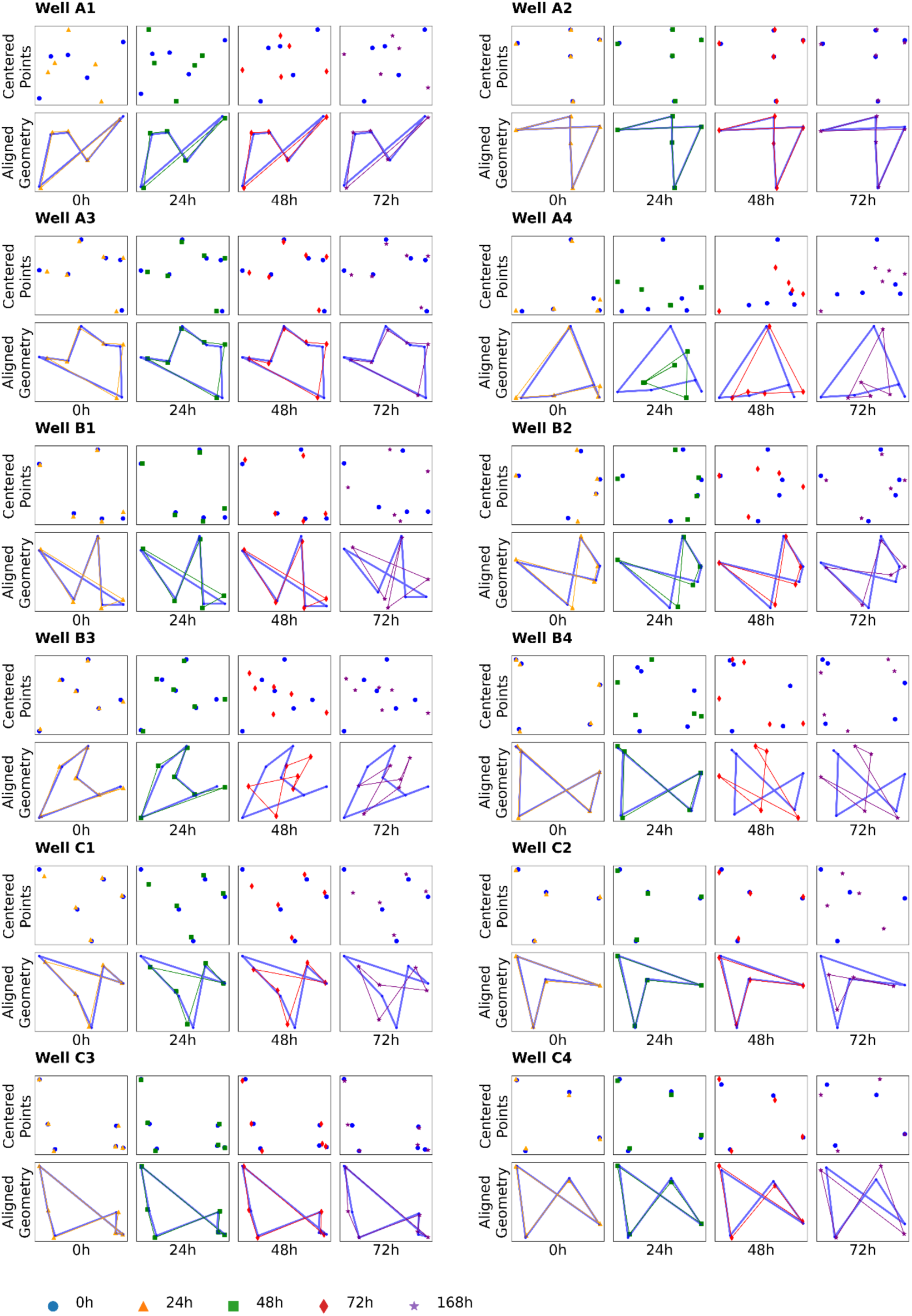
Application of the matching algorithm to the biological data set. The top row of each sub-panel depicts the X and Y coordinates of the spheroids after being centered, and the bottom row depicts the geometry of the points after Procrustes analysis.

**Supplemental Figure 2.**
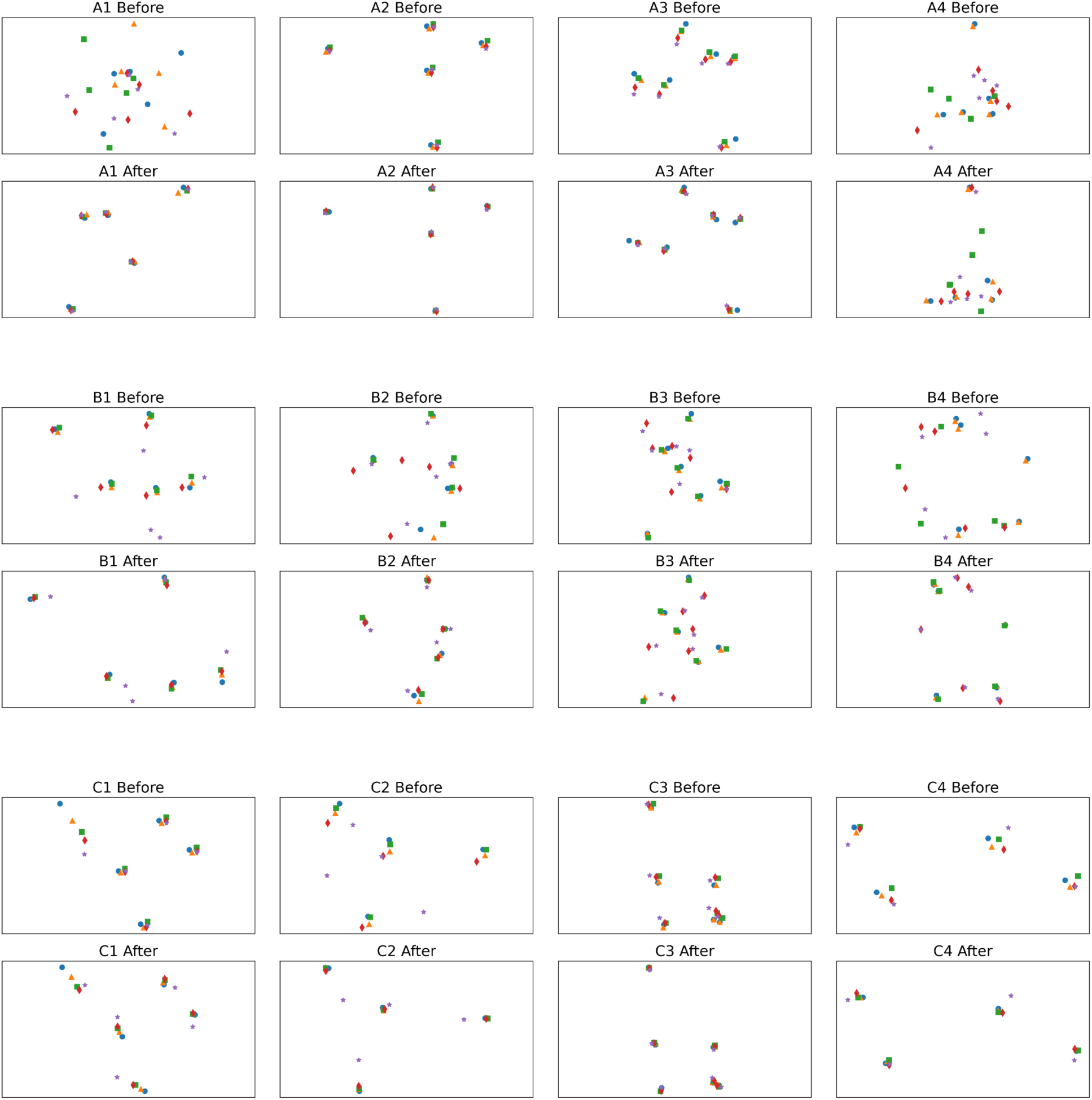
The positions of spheroids before and after applying the matching algorithm.

**Supplemental Figure 3.**
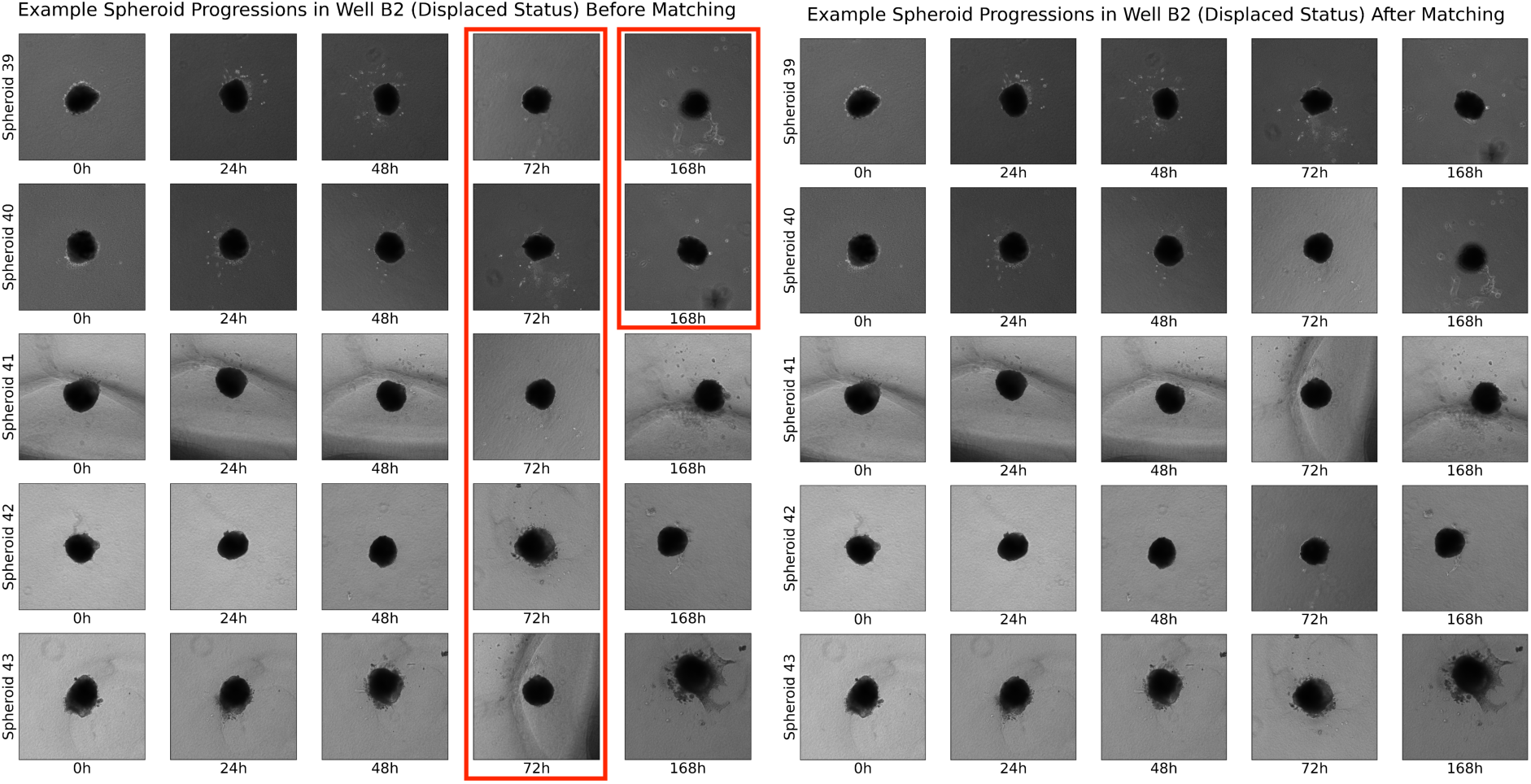
Example application of the algorithm to a well with displaced spheroids in red before (Left) and after matching (Right).

## References

1. Costa, E. C. et al. 3D tumor spheroids: an overview on the tools and techniques used for their analysis. Biotechnol. Adv. 34, 1427–1441 (2016).

2. Thoma, C. R., Zimmermann, M., Agarkova, I., Kelm, J. M. & Krek, W. 3D cell culture systems modeling tumor growth determinants in cancer target discovery. Adv. Drug Deliv. Rev. 69–70, 29–41 (2014).

3. Zhu, Y. et al. 3D Tumor Spheroid and Organoid to Model Tumor Microenvironment for Cancer Immunotherapy. Organoids 1, 149–167 (2022).

4. Book review: *Pharmaceutical* Biotechnology: Drug Discovery and Clinical Applications (2nd Edition). Biotechnol. J. 7, 1061–1062 (2012).

5. Cancer Cell Culture: Methods and Protocols. vol. 2645 (Springer US, New York, NY, 2023).

6. Fevre, R. et al. Combinatorial drug screening on 3D Ewing sarcoma spheroids using droplet-based microfluidics. iScience 26, 106651 (2023).

7. Alicea, G. M. et al. Age-Related Increases in IGFBP2 Increase Melanoma Cell Invasion and Lipid Synthesis. Cancer Res. Commun. 4, 1908–1918 (2024).

8. Bouchard, G. et al. A quantitative spatial cell-cell colocalizations framework enabling comparisons between in vitro assembloids and pathological specimens. Nat. Commun. 16, 1392 (2025).

9. Nazari, S. S. Generation of 3D Tumor Spheroids with Encapsulating Basement Membranes for Invasion Studies. Curr. Protoc. Cell Biol. 87, e105 (2020).

10. Charoen, K. M., Fallica, B., Colson, Y. L., Zaman, M. H. & Grinstaff, M. W. Embedded multicellular spheroids as a biomimetic 3D cancer model for evaluating drug and drug-device combinations. Biomaterials 35, 2264–2271 (2014).

11. Genenger, B., McAlary, L., Perry, J. R., Ashford, B. & Ranson, M. Protocol for the generation and automated confocal imaging of whole multi-cellular tumor spheroids. STAR Protoc. 4, 102331 (2023).

12. Hou, Y., Konen, J., Brat, D. J., Marcus, A. I. & Cooper, L. A. D. TASI: A software tool for spatial-temporal quantification of tumor spheroid dynamics. Sci. Rep. 8, 7248 (2018).

13. Jeong, Y. J. et al. Morphology-guided transcriptomic analysis of human pancreatic cancer organoids reveals microenvironmental signals that enhance invasion. J. Clin. Invest. 133, e162054 (2023).

14. Bengio, Y., Courville, A. & Vincent, P. Representation Learning: A Review and New Perspectives. Preprint at http://arxiv.org/abs/1206.5538 (2014).

15. Holden, M. A Review of Geometric Transformations for Nonrigid Body Registration. IEEE Trans. Med. Imaging 27, 111–128 (2008).

16. COMPUTING THE FRÉCHET DISTANCE BETWEEN TWO POLYGONAL CURVES | International Journal of Computational Geometry & Applications. https://www.worldscientific.com/doi/epdf/10.1142/S0218195995000064.

17. Agrawal, A. et al. Stromal cells regulate mechanics of tumour spheroid. Mater. Today Bio 23, 100821 (2023).

18. Spoerri, L., Gunasingh, G. & Haass, N. K. Fluorescence-Based Quantitative and Spatial Analysis of Tumour Spheroids: A Proposed Tool to Predict Patient-Specific Therapy Response. Front. Digit. Health 3, 668390 (2021).

19. Liu, Y. & Yang, C. Computational methods for alignment and integration of spatially resolved transcriptomics data. Comput. Struct. Biotechnol. J. 23, 1094–1105 (2024).

20. Heussner, R. T. et al. COEXIST: Coordinated single-cell integration of serial multiplexed tissue images. BioRxiv Prepr. Serv. Biol. (2024) doi:10.1101/2024.05.05.592573.

21. Tiriac, H. et al. Organoid Profiling Identifies Common Responders to Chemotherapy in Pancreatic Cancer. Cancer Discov. 8, 1112–1129 (2018).

22. Öhlund, D. et al. Distinct populations of inflammatory fibroblasts and myofibroblasts in pancreatic cancer. J. Exp. Med. 214, 579–596 (2017).

23. Jaffee, E. M. et al. Development and characterization of a cytokine-secreting pancreatic adenocarcinoma vaccine from primary tumors for use in clinical trials. Cancer J. Sci. Am. 4, 194–203 (1998).

24. William Layton & Myron Sussman. Numerical Linear Algebra. (2014).

25. Gower, J. C. Generalized procrustes analysis. Psychometrika 40, 33–51 (1975).

26. Chiu, C.-L., Clack, N., & the napari community. napari: a Python Multi-Dimensional Image Viewer Platform for the Research Community. Microsc. Microanal. 28, 1576–1577 (2022).

27. Kirillov, A., et al. Segment Anything. Preprint at 10.48550/arXiv.2304.02643 (2023).

28. Archit, A. et al. Segment Anything for Microscopy. Nat. Methods 22, 579–591 (2025).

29. Figures 1 and 2 made with Biorender.

